# Biomechanical evaluation of a novel orthogonal angle-stable interlocking nail in a canine femur model Novel orthogonal angle-stable interlocking nail

**DOI:** 10.1101/2021.02.26.433020

**Authors:** D. V. F. Lucena, B. W. Minto, T. A. S. S. Rocha, C. A. S. Malta, J. A. S. Galíndez, L. G. G. G. Dias

## Abstract

Interlockings nails (IN) are orthopedic implants with superior mechanical and, potentially, biological qualities. Despite the countless and indisputable advantages of current angle-stable models, there are still limitations for their use in certain scenarios. The objective of the present study was to describe and biomechanically test a new orthogonal angle-stable intramedullary nail model for veterinary use. The proposed orthogonal angle-stable nail has two 3.8-mm threaded cylindrical holes in each of its portions: in the proximal portion, the holes are 11 mm apart; in the distal portion, the penultimate orifice is positioned at 90 degrees in relation to the last one, with a distance of 5.5 mm between them. The novel orthogonal nail (Group 3 – G3) was evaluated and compared biomechanically with the conventional interlocking nail (Group 1 – G1) and the uniplanar angle-stable nail (Group 2 – G2) by means of destructive torsion and axial compression tests. No statistically significant differences were observed in torsion resistance between the groups in the destructive tests. However, statistical differences were found in stiffness values in the compression tests between the orthogonal (G3) and conventional interlocking (G1) nails (p=0.01) and also between the uniplanar (G2) and interlocking (G1) nails (p=0.001). The new orthogonal nail proved to be biomechanically similar to the uniplanar angle-stable model and superior to the conventional nail. This new arrangement of interlocking screws (orthogonal and closer to each other) potentially enables the fixation of small fragments and at the extremities of long bones in dogs. Nonetheless, further clinical studies are necessary to validate such hypotheses.

## Introduction

A wide range of orthopedic implants are available for fixing long bone fractures in companion animals, and many of them show adequate results as long as they have been applied correctly and following the mechanical and biological principles for decision-making and approach [1]. Among them, the intramedullary nail (IN) can be highlighted, which stands out mechanically because it is applied to the neutral axis of force (medullar canal of the bone) and presents significant resistance to flexion [2]. Due to its potentially minimally invasive application, the method also presents relevant advantages from a biological point of view [3].

INs are considered the implant of choice in many types of fractures in humans [4, 5] and they have been progressively gaining space in medicine of small animals [1, 3], especially in the fixation of diaphyseal fractures of the femur, tibia, and humerus [6]. In spite of the qualities and excellent results reported with the use of IINs in dogs and cats [7], some limitations have been found over time, leading to several modifications and improvements since the first description made by Küntscher in 1940 [8] and in models designed and used in subsequent decades [1, 3, 9 – 11]).

The initially described defect of this model was related to the slack between the nail orifice and the interlocking screws when submitted to compression and torsional force [12, 13]. The lack of a rigid interaction between conventional screws and the holes in the nails creates an unstable environment, predisposing the occurrence of complications [1]. In an attempt to mitigate such risk, systems with rigid interactions between the nail orifice and the screw (angle-stable nail) were developed, which present greater stiffness and resistance than conventional models [1, 10, 11, 14].

Despite the countless and indisputable advantages of the available angle-stable models and their use in juxta-articular fractures [1, 3], there are still some limitations for their use in specific clinical settings or in animals that are too heavy [15]. Improvements in this sense have been observed in IIN systems for use in humans, with an emphasis in multiplanar interlocking nails[16 – 18]. However, these systems remain unavailable or have not been sufficiently tested for clinical application in dogs, cats, and other animals. Orthopedic implants applied in multiple planes are used in challenging scenarios, such as, for example, patients who are considerably heavy or robust, large animals, wild animals, and in metaphyseal or epiphyseal fractures with fragments that are too small [19, 20]. The main advantages of this configuration are correlated with increases in the area moment of inertia and the consequent marked expansion of the system’s mechanical resistance, in addition to the potential capacity for fixing fragments to bone extremities with limited area for fixation [21, 22]. Multiplanar external skeletal fixators and double or triple plates are commonly used for these purposes, with excellent results [23]. However, interlocking intramedullary nails for veterinary use, despite some reports [3], were not designed specifically for this function. Improved nail systems for humans present significant variations in the arrangement of interlocking screws: orthogonal, multiplanar, or in a varied angle [24, 25], which can be used on different fractures.

Inspired by the existence of a gap in the arsenal of options for fixing these fractures in veterinary patients, the objective of this study was to describe and biomechanically test a new orthogonal angle-stable interlocking nail model for veterinary use. It is hypothetically believed that this implant is feasible for application in canine bones and is potentially superior regarding mechanical resistance, thus rendering it capable of minimizing interfragmentary movements.

## Material and Methods

All protocols were approved by the Animal Care and Use Committee of the Júlio Mesquita Filho State University (UNESP).

### Substitute bone models

Substitute bone models were designed using a CAD program (SOLIDWORKS^®^ 2016) to replace and standardize the cortical bone to be used as a test specimen. The bone substitutes were manufactured with polylactic acid (PLA) and had specific dimensions to simulate proximal and distal fracture fragments (19 mm external diameter, 5-mm thick corticals, 9 mm medullary canal diameter, and 98 mm of total length of each segment). They also had pre-defined holes for fixing the screws and a square base (20 mm X 20 mm X 20 mm) for fixation to the mechanical testing machines. The spacing between the proximal and distal fragments was 25 mm in order to simulate the gap of a comminuted fracture. The substitute bone models were designed in two parts: one proximal and one distal. Each group had a specific model (Figure 1). After developing the synthetic models with the CAD program, they were saved in STL format and printed three-dimensionally (3D) in polylactic acid (PLA).

**Fig 1:**
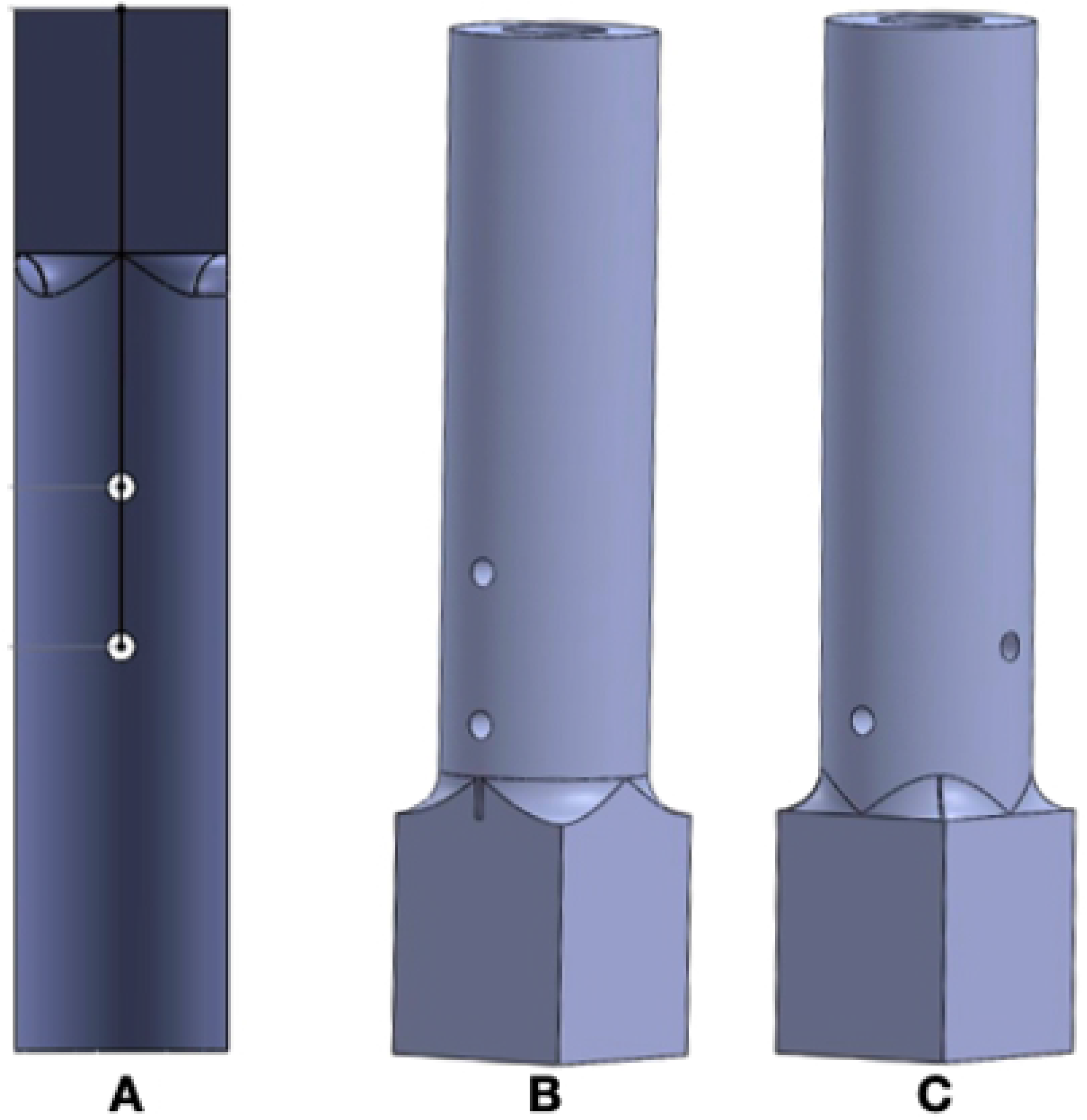
Substitute models of three-dimensionally printed bones with predetermined orifices for the passage of screws. Figure (A) illustrates the proximal model (test specimen) that mimicked the proximal bone fragment (Groups 1, 2, and 3). The model (specimen) shown in Figure (B) was used to mimic the distal bone fragment in Groups 1 and 2. Figure (C) represents the model used in the distal base of Group 3. Note the holes positioned at 90-degree angles from each other. All models had a square base to facilitate their attachment to the universal testing machines.

### Nail models

The nails used in the three studied groups were made of 316L solid stainless steel and measured 8 mm in external diameter and 175 mm in length. The angle-stable nails (Groups 2 and 3) had two threaded cylindrical holes, measuring 3.8 mm in diameter (3.2 mm diameter core) in each portion (proximal and distal). The nails in Group 1 (Control Group), on the other hand, had cylindrical (smooth) holes measuring 3.5 mm in diameter and a 3.0 mm diameter core). The two proximal holes in the nails of all the analyzed groups were 11 mm apart. The distal holes in Groups 1 and 2 were also 11 mm apart (Figure 2A). The orthogonal angle-stable nails (Group 3) presented modifications in the first hole of the distal portion, *i.e*., it was positioned at 90 degrees in relation to the others (two orifices in the proximal portion and the distal orifice in the distal portion); the distance between the two distal holes was 5.5 mm (Figure 2B).

**Fig 2:**
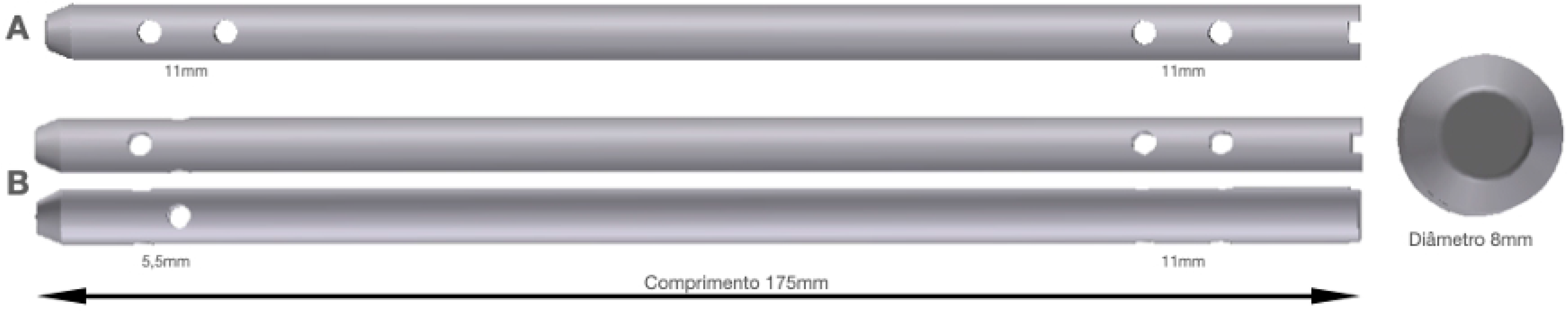
Designs of the angle-stable nails used in the study, measuring a total length of 175 mm and a uniform diameter of 8 mm. Uniplanar nail with 3.8 mm proximal and distal holes distanced 11 mm apart (A). Orthogonal model nail developed for the study, with distal holes measuring 5.5 mm apart and an upper orifice positioned at 90 degrees in relation to the others (B).

In order to perform the mechanical tests, the nails were locked to the models (synthetic test specimens) that mimicked the bone fragments. For the accurate and consistent placement of the locking implants, two pilot holes were made in all the proximal (Groups 1, 2, and 3) and distal (Groups 1 and 2) test specimens, with their centers separated from each other by exactly 11 mm, to coincide with the uniplanar nail holes. Meanwhile, in the distal specimens of Group 3, the two pilot holes (orthogonal) were 5.5 mm apart. The orifices in the synthetic specimens of Groups 2 and 3 measured 2.5 mm in diameter and were previously countersunk (3.2 mm in diameter) regarding the central portion (solid-core) of the screw. Each studied group included 14 test specimens.

### Group configurations

A total of 42 test specimens (comprised of proximal and distal portions, mimicking fractured bone fragments) were used in this study. Each of the three groups had 14 sets of specimens, stabilized with specific metallic implants (intramedullary nails and interlocking implants), denominated constructs. The 14 constructs of each group were divided equally (n=7) to be used in the torsion and axial compression tests. Four bicortical interlocking implants (screws) were used in each construct, two proximally and two distally. All screws were obligatorily fixed to the two corticals of the test specimens. In Group 1, a conventional intramedullary nail was used, *i.e*., without locking between the hole and the screw. In this group, the used cortical type screws measured 3.5 mm in diameter and were arranged on a single plane. In Group 2 (G2), the angle-stable nail model (threaded) was used, as well as uniplanar locking, with screws with an external diameter of 3.8 mm. Finally, in Group 3 (G3), the screws had an outer diameter of 3.8 mm and were arranged on the same plane in the proximal portion and orthogonally in the distal portion of the specimen.

### Biomechanical test

The biomechanical properties of the constructs (synthetic specimens-nail) were evaluated by means of destructive axial compression and torsion tests. The obtained data were recorded in Newtons (N) in the axial compression assessment and Newtons-meter (Nm) in the torsion test

For the compression tests, a universal testing machine (UTM – model DL 10000) was used, with a standardized displacement of 0.05 mm per second. The tested constructs were positioned in the upright position so that the machine could exert an eccentric force of compression in relation to the specimen, simulating the main load applied to appendicular bones during locomotion (Figure 3).

**Fig 3:**
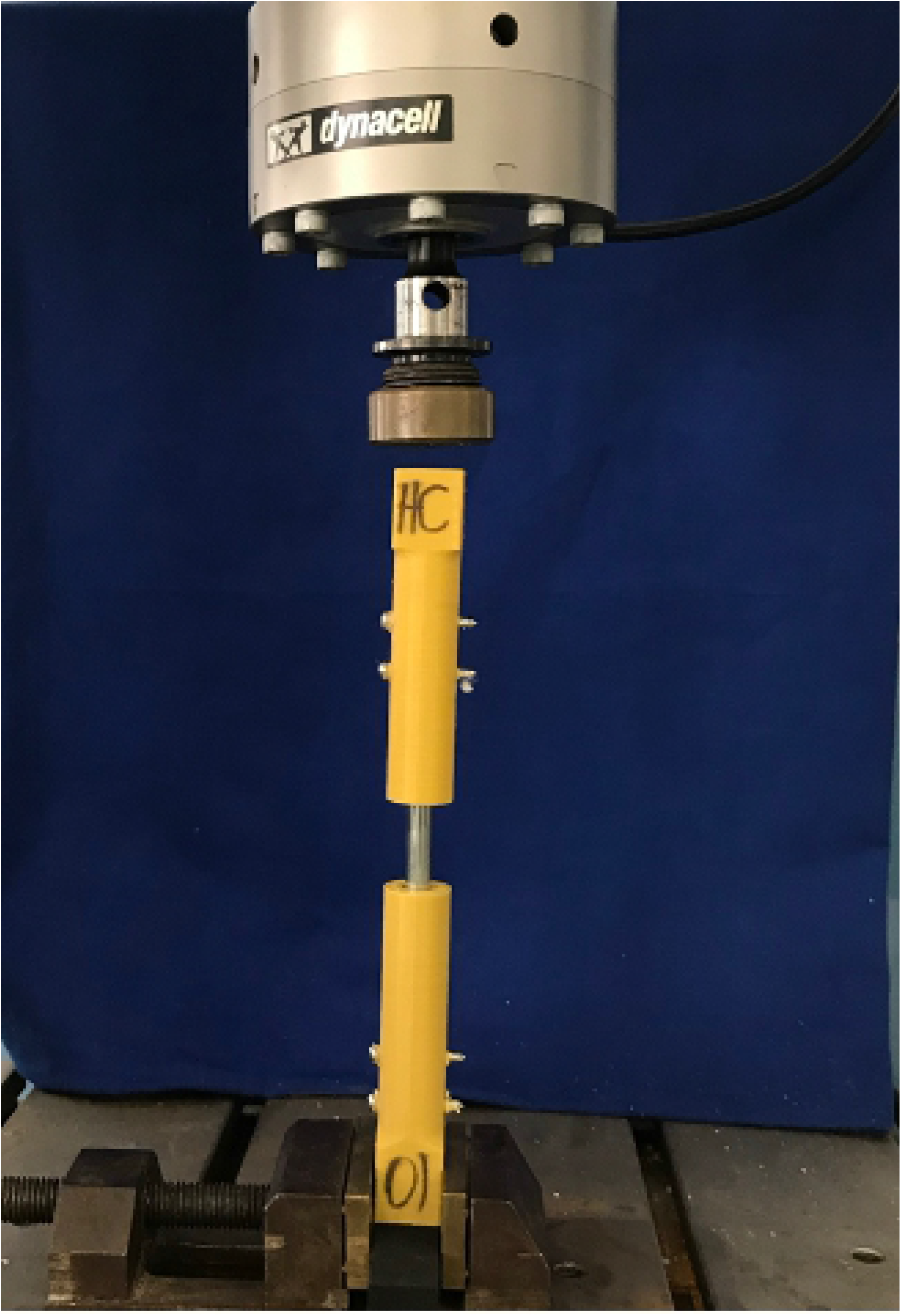
Photographic image of the construct positioned vertically within the servo-hydraulic testing machine. The solid block on the distal portion of the specimen was fixed to a specific device of the testing machine, remaining immobile. The load cell exerted force at the base of the proximal portion of the test specimen. When the test started, the proximal portion of the specimen was pressed towards its distal portion.

**Fig 4:**
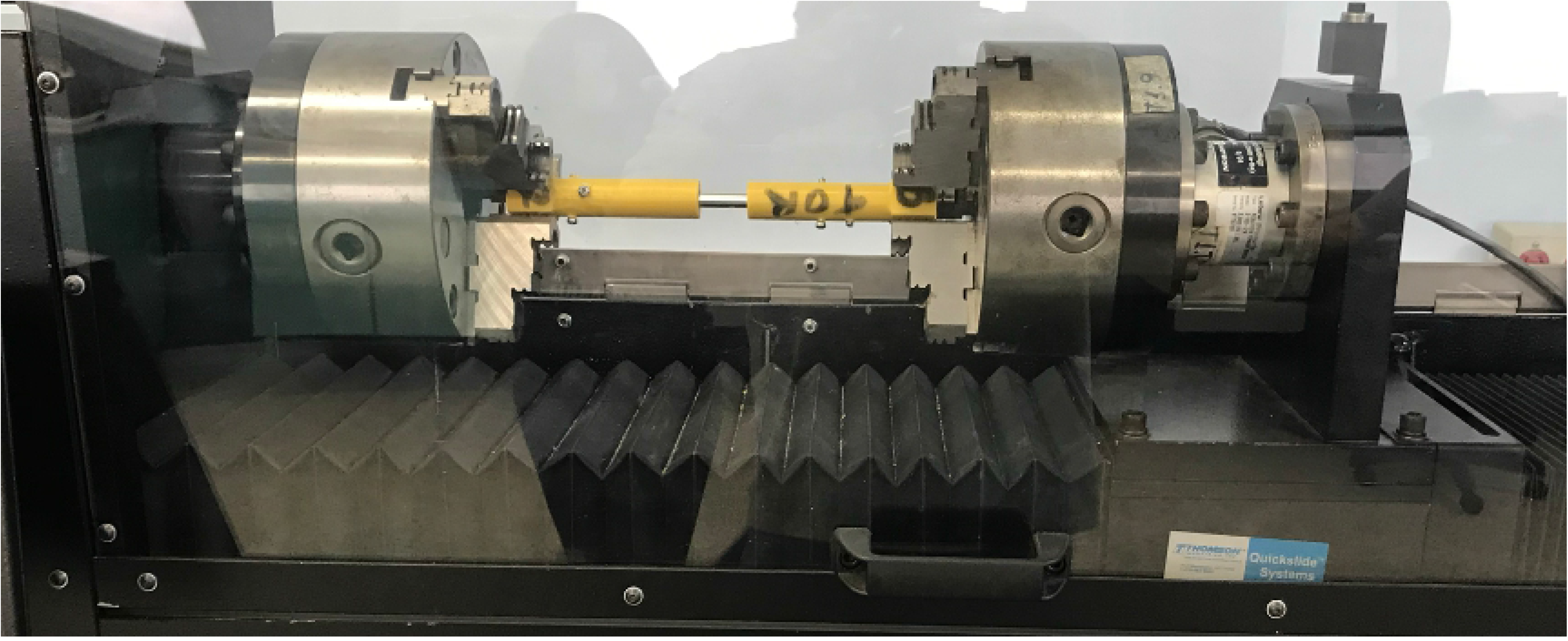
Photographic image of the construct coupled to the servo-hydraulic testing machine, which was attached by two pulleys to the square base of the synthetic bone model. When the test started, the pulley located on the right in the picture was responsible for promoting rotation.

The torsion tests were carried out using the Instron torsion testing machine (model 55MT), which consisted of two pulleys that enabled the fixation of the constructs in horizontal position. One side of the construct was maintained stable (stationary), while the other side (proximal portion) was subjected to torsion. The three groups were loaded until failure, with a displacement rate of one degree per second.

### Statistical analysis

All statistical procedures were conducted with the SPSS software, version 20.0, for Windows. The analysis of the normal distribution of data was performed using the Shapiro-Wilk test, which resulted in non-parametric distribution. Three variables were evaluated in this study: the peak torque when torsion was carried out, stiffness, and the maximum force registered during the compression tests. All variables were analyzed using the Kruskal-Wallis test, in addition to Dunn’s *post-hoc* test, performed in order to establish the p-value between the G1, G2, and G3 nail groups. A 5% (p<0.05) level of significance was adopted in all the statistical analyses.

## Results

Among the three groups, G2 obtained the highest maximum torque value (maximum force of 29.47 Nm), as well as the highest mean (23.39 Nm), in the torsion test. During the assessment, the specimens of this group exhibited plastic deformation when reaching the maximum load. Since it was impossible to detect deformation or breakage of the screws or nails, the failure occurred in the synthetic specimens themselves. In G3, the maximum recorded force was 27.16 Nm, with a mean of 21.76 Nm. Similar to G2, the plastic deformation occurred only in the synthetic specimens, without macroscopic alterations in the screws or nails. As for G1, the maximum and mean force values of this group were 24.72 Nm and 21.27 Nm, respectively, representing the lowest values found in the three groups. After the end of the test, it was possible to observe plastic deformations in the screws in that group. As in the other groups, the test was also interrupted when the synthetic specimen displayed failure.

In the compression test, two factors were evaluated: the stiffness of the constructs and the maximum force they sustained. The maximum and mean resistance observed in G1 corresponded to 12,128 N and 11,254 N, respectively. In G2, the maximum and mean resistance values were 12,944 N and 11,648 N, respectively. G3 obtained the highest values of maximum force, reaching 13,053 N in maximum resistance.

The stiffness values found in G1 were 1,873 N of mean stiffness and a standard deviation of 153.1 N. In G2, a mean stiffness of 2,522 N was observed, as well as a standard deviation of 153.1 N. Meanwhile, in G3, the mean value found was 2,466 N, with a standard deviation of 95.94 N.

At the end of the compression tests, all constructs underwent macroscopic evaluation to identify possible plastic deformations in the screws. In G1, all constructs showed evident screw deformation, and in some specimens, the deformation was such that the ends of the screws (head and tip) entered the medullary cavity of the synthetic specimens. In G2, all specimens underwent plastic deformation in the distal screws, although to a lesser extent when compared to G1. The same alteration in G2 was observed in three specimens from G3. One of the constructs exhibited nail deformation (flexion), although the screws did not show any apparent alterations.

According to the obtained results, it can be noted that only the stiffness values were statistically significant (Kruskal-Wallis p-value of p<0.001). Dunn’s *post-hoc* test revealed a significant difference between the orthogonal and interlocking nail (p=0.01) and also between the uniplanar and interlocking nail (p=0.001). The other analyses were not statistically different; their p-values were: peak torque (p=0.339) and maximum force (p=0.739) (Figure 5).

**Fig 5:**
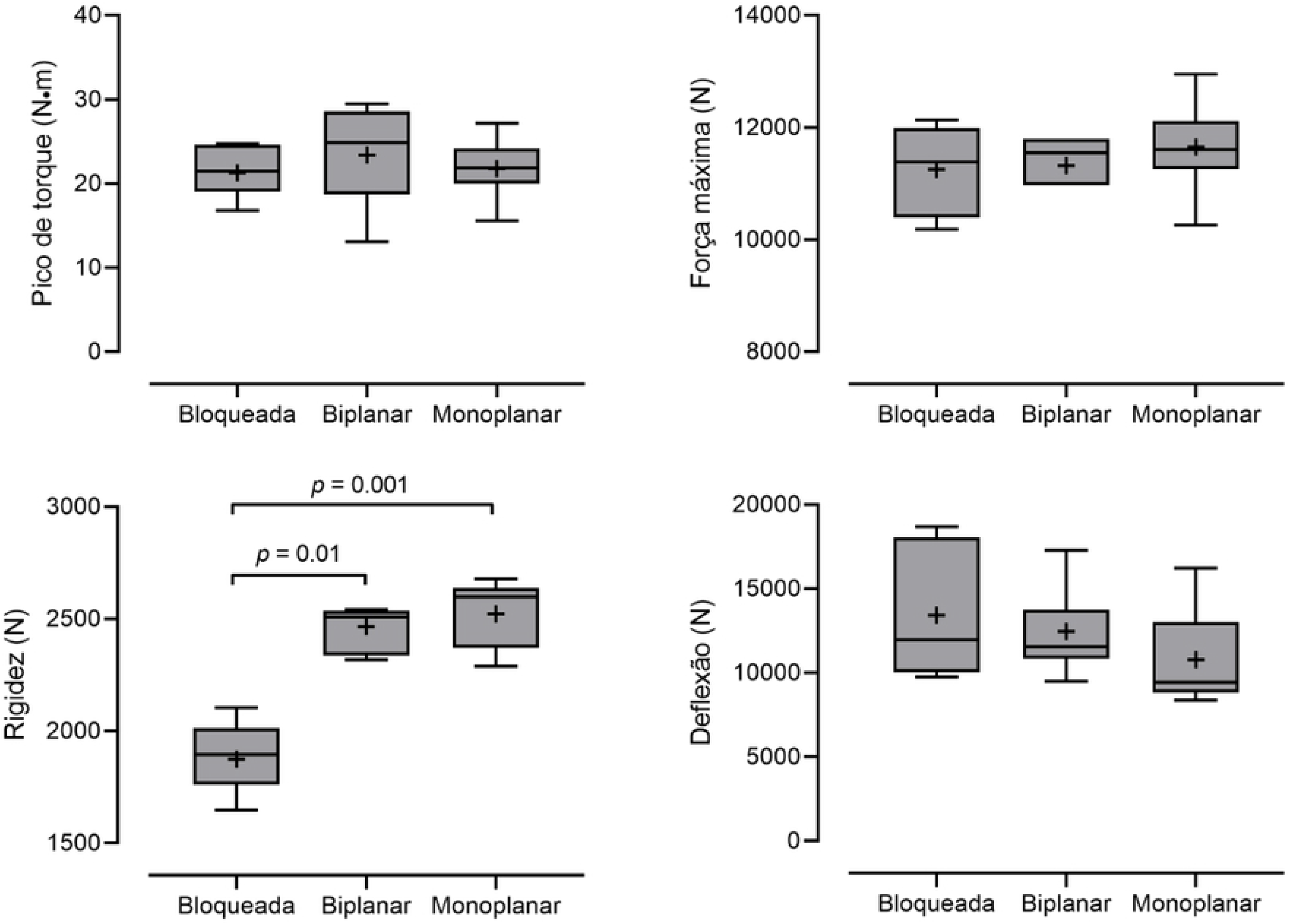
Image showing the graphs in Boxplot format. The mean and median are represented by the plus symbol (+) and the line inside the boxes, respectively. Lines above the boxes indicate the maximum values, while those below them indicate the minimum values.

## Discussion

A vast arsenal of techniques and implants are currently available to veterinary surgeons, allowing them to make more effective and appropriate decisions in various scenarios. Despite this reality, certain fractures remain major challenges, and the complication rates are higher than expected [26]. Periarticular fractures, with tiny fragments, or those in patients who are too heavy or large, are noteworthy since they still challenge surgeons and the most modern implants [27]. The present study demonstrated the mechanical characteristics of a new orthogonal angle-stable interlocking intramedullary nail implant (Group 3) and compared it with models classically used in veterinary patients. The initial hypothesis that it could exhibit good biomechanical behavior and be superior to the tested models was not entirely confirmed, given that the new device significantly increases the stiffness of the construct under axial load. However, it was not significantly superior to Group 2 in the test. In addition, the change in the positioning of one of the screws, which created a right angle between the two interlocking implants in the distal specimen, combined with the more distal location of these two implants on the nail, significantly increased the stiffness of the construct under axial load and potentially allowed its fixation to fragments closer to the adjacent joint in a hypothetical clinical scenario. The hypothesis raised herein has yet to be evaluated in an *in vivo* study.

The torsion tests applied to the constructs in Groups 2 and 3 were interrupted at the time of failure of the synthetic specimens, with a load of 2,466 Nm and 2,522 Nm, respectively. In contrast, some sets from G1 exhibited plastic deformation of the screws before specimen failure and with less load (1,873 Nm), indicating this group’s inferiority in relation to the others regarding resistance to the applied torsional loads, despite their being undoubtedly supraphysiological [28]. The differences between the internal diameters of the screws/bolts used in Groups 2 and 3 (3.2 mm) compared to Group 1 (2.4 mm) infer a marked increase in the area moment of inertia (AMI) of the screws, justifying such an event [28]. Plenert [29] demonstrated greater resistance of the sets formed by nails interlocked with screws of increased diameter in compression and torsion tests. Despite the methodological differences, both studies used supraphysiological loads, which does not invalidate the sets’ ability to be resistant enough in a hypothetical clinical scenario.

In the absence of statistically significant differences between the maximum loads in the destructive mechanical torsion test between Groups 2 and 3, the initial hypothesis that the orthogonal nail would be superior also in the torsion tests was annulled. It has already been proven that angle-stable models have superior resistance in mechanical tests when compared to traditional models since they promote greater stability [10, 11], a fact that, once again, was made evident in our study. However, despite the orthogonal model having shown lower values than the uniplanar model, these values are representative. We believe that this model also has sufficient torsional resistance to withstand physiological loads applied by dogs during locomotion.

The addition of oblique screws significantly increases the mechanical stability of constructs with intramedullary nails [16]. The authors of this study believe that the role of the orthogonal screw will be fundamental in metaphyseal fractures since it maintains axial alignment due to the absence of contact between the nail and the cortical bone. The cross-sectional geometry of oblique screws also adds stiffness to the construct, opposing torsional movement symmetrically around the longitudinal axis [28]The authors of the present study anticipate that future cyclic tests with this nail model will help to better understand the mechanical behavior of this type of implant.

Orthogonal fixation is often used in other fixation methods, such as external skeletal fixators [30]. Pins positioned at 60 or 90 degrees of angulation enable an increase in resistance, thus improving stiffness against torsional deformation [31]. Favorable mechanical and clinical results with the use of circular fixators using orthogonal pins have brought popularity to this stabilization method that is widely used in veterinary medicine to correct several types of fractures [32, 33]. In the same way that satisfactory results have emerged with the use of orthogonal fixators, the authors of the present study believe that they may be similar to orthogonal intramedullary nails.

A material’s stiffness is directly influenced by the load volume necessary to promote its deformation [28]. The stiffness recorded in the axial compression test in the orthogonal nail group was superior to the other groups and statistically relevant in relation to the interlocking nails (G1) (p = 0.01). Even though there were no statistically significant differences between the groups of angle-stable intramedullary nails, the values were numerically higher in G3. We believe that, given that these two groups used screws with similar diameters, orthogonal positioning may have potentially influenced the results. Implants positioned to allow the orthogonal placement of their screws have proven to be mechanically superior in other studies involving several types of implants [19, 20]. In the study by Baseri [34] where the authors compared the use of angle-stable nails and locking plates in the treatment of distal tibial fractures in humans, the authors stated that nails are mechanically more efficient than locking plates; however, the use of uniplanar nails requires the transverse fixation of Poller screws to avoid shear. Since orthogonal nails used to stabilize proximal or distal metaphyseal fractures in humans promote fixation in multiple planes, this method has shown to be superior to other models [18]. Their surgical applicability confirms the results obtained in mechanical tests, and minimizes complications with the use of fixed uniplanar mid-lateral screws that can result in nail translation, causing misalignment in bone valgus or varus [].

Although the initial hypothesis that the orthogonal angle-stable nail model would improve mechanical performance has not yet been statistically proven, we can state that the new nail model represents a construct capable of resisting supraphysiological loads, similarly to other models already tested and considered excellent from a mechanical point of view. The preliminary results shown in this study indicate that the influence of angular stability seems to be stronger when compared to uniplanar stability in axial loads. The present study, being the first to test an orthogonal nail model for veterinary patients, opted for the destructive test in a single cycle in order to confirm the construct’s ability to withstand supraphysiological loads and identify possible moments of failure. Further studies testing the behavior of this nail model in cyclic trials and clinical tests are essential to better evaluate the real advantages of orthogonal interlocking nails.

## Limitations

The present study evaluated two loads in a single destructive cycle. Although the two tested loads were the primary ones in the intramedullary nail assessment, it is essential that the results recorded herein are associated with cyclic biomechanical testing, the best method to evaluate the behavior of the construct in patients *in vivo*, since it is known that fatigue-related failure is the main type of failure reported in orthopedic implants.

## References

[1] Déjardin L, Guiot L, Pfeil D. Interlocking nails and minimally invasive osteosynthesis. Veterinary clinical small animal. 2012; (42):935–962.

[2] Goett S, Sinnott M, Ting D, Basinger R, Haut R, Déjardin L. Mechanical Comparison of an Interlocking Nail Locked with Conventional Bolts to Extended Bolts Connected with a Type-Ia External Skeletal Fixator in a Tibial Fracture Model. Veterinary Surgery. 2007; (36): 279–286.

[3] Déjardin L, Perry K, Pfeil D, Guiot L. Interlocking nails and minimally invasive osteosynthesis. Veterinary clinical small animal. 2019; (50):67–100.

[4] Chaudhary P, Maharjan R, Kalawar R, Baral P, Shah A. Randomized controlled trial comparing open versus closed interlocking nail for closed fracture shaft of femur in Adults. International Journal of Orthopaedics Sciences. 2017, (1): 591–595.

[5] Deepak C, Chethan B. A study of functional outcome of femoral diaphyseal fractures by closed reduction and internal fixation using intramedullary interlocking nail in adults. International Journal of Orthopaedics Sciences. 2019, (1): 132–138.

[6] Alt V, Leung K, Kempf I, Haarman H, Taglang G, Seidel H, et al. Practice of intramedullary locked nails: New developments in techniques and applications. Berlin: Springer-Verlag; 2006.

[7] Durall I, Diaz M. Early experience with the use of an interlocking nail for the repair of canine femoral shaft fractures. Veterinary Surgery. 1996; 25(5):397–406.

[8] Dueland R, Berglund L, Vanderby R, Chao E. Structural properties of interlocking nails, canine femora, and femur-interlocking nail constructs. Veterinary Surgery. 1996; 25(5):386–396.

[9] Bhat S, Aithal H, Kinjavdekar P, Amarpal, Zama M, Gope P, Pawde A, Ahmad R, Gugjoo M. Na in vivo Biomechanical investigation of na interlocking nail system developed for buffalo tibia. Vet Comp Orthop Traumatol. 2014; (27): 36 – 44.

[10] Déjardin L, Lansdowne J, Sinnott M, Sidebotham C, Haut R. In vitro mechanical evaluation of torsional loading in simulated canine tibiae for a novel hourglass-shaped interlocking nail with a self-tapping tapered locking design. AJVR. 2006; (67): 678 – 685.

[11] Déjardin L, Guillou R, Ting D, Sinnott M, Meyer E, Haut R. Effect of bending direction on the mechanical behavior of interlocking nail systems. Vet Comp Orthop Traumatol. 2009; (22): 264 – 269.

[12] Duhautois B. Use of veterinary interlocking nails for diaphyseal fractures in dogs and cats: 121 cases. Vet Surg. 2003; 32: 8–20.

[13] Moses P, Lewis D, Lanz O, Stubbs W, Cross A, Smith K. Intramedullary interlocking nail stabilisation of 21 humeral fractures in 19 dogs and one cat. Australian veterinary journal. 2002; 80(6):336–43.

[14] Déjardin L, Cabassu J, Guillou R, Villwock M, Guiot L, Haut R. In Vivo Biomechanical Evaluation of a Novel Angle‐Stable Interlocking Nail Design in a Canine Tibial Fracture Model. Vet Surg. 2014; 43: 271–281.

[15] DeTora M, Boudrieau R. Complex angular and torsional deformities (distal femoral malunions). Vet Comp Orthop Traumatol. 2016; 29: 416–425.

[16] Laflamme G, Heimlich D, Stephen D, Kreder H, Whyne C. Proximal Tibial Fracture Stability with Intramedullary Nail Fixation Using Oblique Interlocking Screws. Journal of Orthopedic Trauma. 2003; 17: 496 – 502.

[17] Lenz M, Gueorguiev B, Richards R, Mückley T, Hofmann G, Höntzsch D, Windolf M. Fatigue performance of angle-stable tibial nail interlocking screws. International Orthopaedics. 2013; 37: 113–118.

[18] F. Hoegel, S. Hoffmann, P. Weninger, V. Bühren. Biomechanical comparison of locked plate osteosynthesis, reamed and unreamed nailing in conventional interlocking technique, and unreamed angle stable nailing in distal tibia fractures. Journal of Trauma and Acute Care Surgery. 2012; 73: 933 – 938.

[19] Brown G, Kalff S, Gemmill T, Pink J, Oxley B, McKee W, Clarke S. Highly Comminuted, Articular Fractures of the Distal Antebrachium Managed by Pancarpal Arthrodesis in 8 Dogs. Vet Surg. 2016; 45: 44–51.

[20] Schmierer P, Smolders L, Zderic I, Gueorguiev B, Pozzi A, Knell S. Biomechanical properties of plate constructs for feline ilial fracture gap stabilization. Vet Surg. 2018; 1–8.

[21] Kosmopoulos V, Nana A. Dual Plating of Humeral Shaft Fractures: Orthogonal Plates Biomechanically Outperform Side-by-Side Plates. Clin Orthop Relat Res. 2014; 472 (4): 1310–7.

[22] Cheng T, Xia R, Yan X, Luo C. Double-plating fixation of comminuted femoral shaft fractures with concomitant thoracic trauma. Journal of International Medical Research. 2017; 0(0): 1–8.

[23] Tian D, Jing J, Qian J, Li J. Comparison of two different double-plate fixation methods with olecranon osteotomy for intercondylar fractures of the distal humeri of young adults. Experimental and therapeutic medicine. 2013; 6: 147–151.

[24] Argintar E, Cohen M, Eglseder A, Edwards S. Clinical Results of Olecranon Fractures Treated With Multiplanar Locked Intramedullary Nailing. J Orthop Trauma. 2013; 27: 140–144

[25] Shani R, Morris R, Gugala Z, Lindsey R. Biomechanical Properties of Conventional Versus Angular Stabilized-Intramedullary Nail Distal Interlocking Screw Configurations in a Distal Tibia Fracture Model. TOJ. 2015; 1: 69–77.

[26] Vallefuoco R, Pommellet H, Savin A, Decambron A, Manassero M, Viateau V, Gauthier O, Fayolle P. Complications of appendicular fracture repair in cats and small dogs using locking compression plates. Vet Comp Orthop Traumatol. 2016; 25: 46 – 52.

[27] Moffatt F, Kulendra K, Meeson R. Repair of Y-T Humeral Condyle Fractures with Locking Compression Plate Fixation. Vet Comp Orthop Traumatol 2019;32:401–407.

[28] Chao P. Lewis D, Kowaleski M, Pozzi A. Biomechanical Concepts Applicable to Minimally Invasive Fracture Repair in Small Animals. Vet Clin Small Anim. 2012; 42: 853–872.

[29] Plenert T, Garlichs G, Nolte I. Harder L, Hootak M, Kramer S, Behrens B, Bach J. Biomechanical comparison of a new expandable intramedullary nail and conventional intramedullary nails for femoral osteosynthesis in dogs. Plos One. 2020; 1 – 20.

[30] Lenarz C, Bledsoe G, Watson J. Circular External Fixation Frames with Divergent Half Pins A Pilot Biomechanical Study. Clin Orthop Relat Res. 2008; 466(12): 2933–2939.

[31] Roberts S, Dodds J, Perry K, Beck D, Seligson D, Voor M. Hybrid External Fixation of the Proximal Tibia: Strategies to Improve Frame Stability. Journal of Orthopaedic Trauma. 2003; 17: 415 – 420.

[32] Yilmaz E, Belhan O, Karakurt L, Arslan N, Serin. E. Mechanical performance of hybrid Ilizarov external fixator in comparison with Ilizarov circular external fixator, Clinical Biomechanics. 2003; 18: 518 – 522.

[33] Spcoe M, Rovesti G, Griffon D, Elkhatib O, Mudrock R, Kurath P. Biomechanical comparison of strategies to adjust axial stiffness of a hybrid fixator, Veterinary Comparative Orthopedics Traumatologic. 2012; 25: 224 – 230.

[34] Baseri A, Bagheri M, Rouhi G, Aghighi M, Bagheri N. 4 Fixation of distal tibia fracture through plating, nailing, and nailing with Poller screws: A comparative biomechanical-based experimental and numerical investigation. Journal of Engineering in Medicine. 2020; 10: 1 – 10.

[35] Singh K, Singh Y, Singh P, Goyal R, Chandra H. Unreamed intramedullary nailing with oblique proximal and biplanar distal interlocking screws for proximal third tibial fractures. Journal of Orthopedic Surgery. 2009; 17: 23 – 27.

